# Oxytocin signaling in the posterior hypothalamus prevents hyperphagic obesity in mice

**DOI:** 10.1101/2021.11.29.470473

**Authors:** Kengo Inada, Kazuko Tsujimoto, Masahide Yoshida, Katsuhiko Nishimori, Kazunari Miyamichi

**Author notes:** Correspondence (K.I.), (K.M.).

## Abstract

Decades of studies have revealed molecular and neural circuit bases for body weight homeostasis. Neural hormone oxytocin (OT) has received attention in this context because it is produced by neurons in the paraventricular hypothalamic nucleus (PVH), a known output center of hypothalamic regulation of appetite. OT has an anorexigenic effect, as shown in human studies, and can mediate satiety signals in rodents. However, the function of OT signaling in the physiological regulation of appetite has remained in question, because whole-body knockout (KO) of *OT* or *OT receptor (OTR)* has little effect on food intake. We herein show that acute conditional KO (cKO) of OT selectively in the adult PVH, but not in the supraoptic nucleus, markedly increases body weight and food intake, with an elevated level of plasma triglyceride and leptin. Intraperitoneal administration of OT rescues the hyperphagic phenotype of the PVH OT cKO model. Furthermore, we show that cKO of OTR selectively in the posterior hypothalamic regions, especially the arcuate hypothalamic nucleus, a primary center for appetite regulations, phenocopies hyperphagic obesity. Collectively, these data reveal that OT signaling in the arcuate nucleus suppresses excessive food intake.

## Introduction

Appetite is one of the strongest desires in animals. The consumption of nutritious foods is a primitive pleasure for animals because it is essential for survival. Yet, excessive food intake leads to obesity and increases the risk of disease. Understanding the neurobiological bases of appetite regulation is therefore an urgent issue, given that the body mass index of humans has increased dramatically over the last 40 years (NCD Risk Factor Collaboration, 2016).

Decades of studies in rodents have revealed molecular and neural circuit bases for body weight homeostasis (Andermann and Lowell, 2017; Sternson and Eiselt, 2017; Sutton et al., 2016). Classical studies with mechanical or electrical lesioning, as well as recent molecular or genetic dissections, both support the critical roles of appetite regulation by neurons in the arcuate hypothalamic nucleus (ARH), in particular, those expressing orexigenic agouti-related protein (Agrp) and anorexigenic pre-opiomelanocortin (Pomc) (Choi et al., 1999; Ollmann et al., 1997). These neurons receive both direct humoral inputs and neural inputs of interoception (Bai et al., 2019) to regulate food intake antagonistically at various time scales (Krashes et al., 2014; Sternson and Eiselt, 2017). The paraventricular hypothalamic nucleus (PVH) is one of the critical output structures of the primary appetite-regulating ARH neurons. Silencing PVH neurons phenocopies the overeating effect observed in the activation of Agrp neurons, whereas activating PVH neurons ameliorates the overeating caused by the acute activation of Agrp neurons (Atasoy et al., 2012; Garfield et al., 2015). Melanocortin-4 receptor (MC4R)-expressing neurons in the PVH are the key target of ARH Agrp and Pomc neurons. PVH MC4R neurons are activated by α-melanocyte-stimulating hormone provided by Pomc neurons and inhibited by GABAergic Agrp neurons, and are supposed to transmit signals to the downstream target regions in the midbrain and pons (Garfield et al., 2015; Stachniak et al., 2014; Sutton et al., 2016).

Although PVH MC4R neurons have been relatively well documented (Balthasar et al., 2005; Garfield et al., 2015), other PVH cell types may also mediate output signals to control feeding and energy expenditure (Sutton et al., 2016). However, little is known about the organization, cell types, and neurotransmitters by which appetite-regulating signals are conveyed to other brain regions. Neural hormone oxytocin (OT), which marks one of the major cell types in the PVH, has received attention in this context (Leng and Sabatier, 2017; Onaka and Takayanagi, 2019). The anorexigenic effect of OT has been shown in humans (Lawson et al., 2015; Thienel et al., 2016), and genetic variations of *OT receptor(OTR)* have been implicated as a risk factor of overeating (Catli et al., 2021; Davis et al., 2017). In rodents, OT administration has been shown to suppress increases in food intake and body weight (Maejima et al., 2018). Pons-projecting OT neurons have been shown to be active following leptin administration (Blevins et al., 2004), and knockdown of OTR in the nucleus of the solitary tract has been reported to alter feeding patterns (Ong et al., 2017). In addition, *OTR*-expressing neurons in the ARH have been shown to evoke acute appetite-suppression signals when opto- or chemo-genetically activated (Fenselau et al., 2017). Despite the importance of the OT-OTR system in the food intake and homeostasis of body weight (Maejima et al., 2018), knockout (KO) and ablation studies still question such findings (Sutton et al., 2016; Worth and Luckman, 2021). For example, *OT* or *OTR* KO mice showed increased body weight at around 4 months old (termed late-onset obesity), while their food intake was not different from that of wild-type mice (Camerino, 2009; Takayanagi et al., 2008). Diphtheria toxin-based genetic ablation of *OT*-expressing cells in adult mice increased the body weight of male mice with a high-fat diet, but not those with normal chow, and in both cases, food intake was unaffected (Wu et al., 2012b). To revisit the function and sites of action of OT signaling in the regulation of feeding, acute conditional KO (cKO) mouse models would be useful.

Here, we describe OT cKO phenotypes related to hyperphagic obesity. Our approach offers the following two advantages over previous studies: (i) the OT gene can be knocked out in adult mice, avoiding the influence of possible developmental and genetic compensations (El-Brolosy et al., 2019); and (ii) the manipulation can be restricted to the brain, or even to a single hypothalamic nucleus, providing a resolution that exceeds previous studies. Owing to these advantages, we show that OT cKO increases both body weight and food intake. The suppression of overeating and overweighting is predominantly regulated by OT neurons in the PVH, leaving OT neurons in the supraoptic nucleus (SO) with only a minor role. We further show that OTR-expressing neurons in the posterior part of the hypothalamus, especially the ARH, mediate the overeating-suppression signals generated by OT neurons.

## Results

### Conditional KO of PVH *OT* increases body weight and food intake

To examine the necessity of OT for the regulation of food intake, we prepared recently validated *OT^flox/flox^* mice (Inada et al., 2022). In this line, Cre expression deletes floxed exon 1 of the *OT* gene, resulting in the loss of transcription of *OT* mRNA (Inada et al., 2022). To perform the cKO in PVH OT neurons, we first crossed *OT^flox/flox^* and *OT* KO (*OT*^-/-^) mice and obtained *OT^flox/-^* mice. Then, we injected *AAV-Cre* into the bilateral PVH of 8-week-old *OT^flox/-^* male mice (Figure 1A and 1B). The number of neurons expressing *OT*, visualized by *in situ* hybridization (ISH), significantly decreased within 3 weeks after the *AAV-Cre* injection (Figure 1C). The body weight of *OT^flox/-^* mice that received *AAV-Cre* injection started to deviate from the controls at around 3 weeks after the injection (Figure 1D). At 5 weeks after the injection, we compared the body weight of *OT^flox/-^* mice that received *AAV-Cre* injection with the wild-type (*OT*^+/+^), *OT^-/-^*, and *OT^flox/-^* mice that received vehicle injection. We found that *AAV*-*Cre*-injected *OT^flox/-^* mice were heavier than those in the other groups (Figure 1E and 1F). Importantly, this increase in body weight was considered unlikely to be a reflection of late-onset obesity, as previously reported (Camerino, 2009; Takayanagi et al., 2008), because we did not find a significant difference between the wild-type and *OT^-/-^* mice (Figure 1F). Next, we analyzed the relationship between the number of remaining *OT*+ neurons and body weight. We found that mice with a fewer number of remaining *OT*+ neurons showed a heavier body weight (Figure 1G). We also found an increase in food intake: the daily food intake of *OT^flox/-^* mice that received *AAV-Cre* injection was significantly larger at >4 weeks after the injection (Figure 1H), and the total food intake during the 5 weeks after the injection was also larger in the mice that received *AAV-Cre* injection (Figure 1I). Of note, these effects are not due to the nonspecific toxicity of *AAV-Cre* injection per se, as *AAV-Cre* injection to wild-type mice did not alter body weight or daily food intake (Figure S1A–D). These results demonstrate that the cKO of *OT* evokes increases in both body weight and food intake.

**Figure 1.**
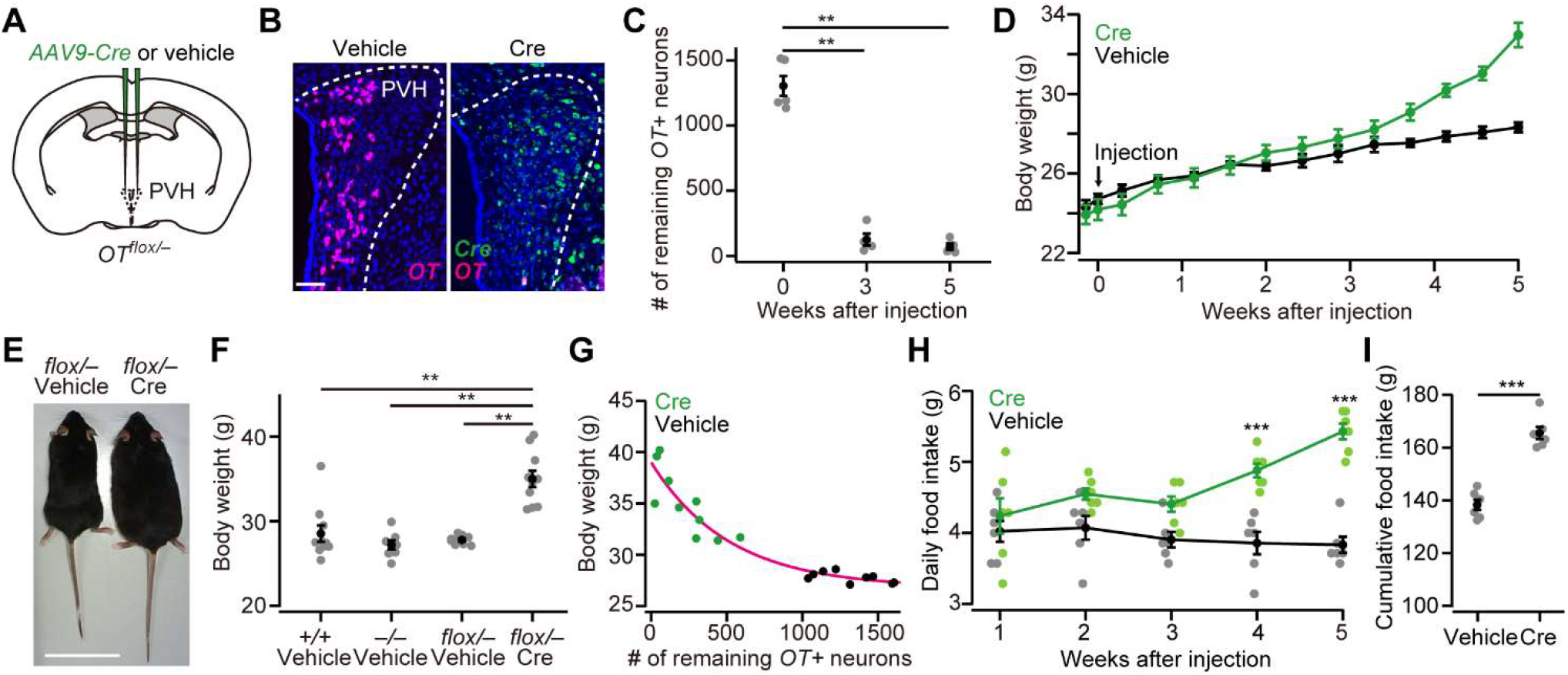
OT cKO in PVH induces an increase in body weight and food intake. (A) Schematic of the virus injection. *AAV-Cre* or vehicle was injected into the bilateral PVH of *OT^flox/-^* male mice. (B) Representative coronal sections of left PVH from *OT^flox/-^* mice received vehicle (left) or *AAV-Cre* (right) injection. Data were obtained at 5 weeks after the injection. Magenta and green represent *OT* and *Cre in situ* stainings, respectively. Blue, DAPI. Scale bar, 50 μm. (C) The number of remaining *OT*+ neurons in the PVH of mice that received *AAV-Cre* injection. **p < 0.01, one-way ANOVA with post hoc Tukey’s HSD. N = 5 each. (D) Time course of body weight after *AAV-Cre* or vehicle injection. N = 6 each. (E) Representative photos of *OT^flox/-^* mice that received either vehicle (left) or *AAV-Cre* injection (right). Five weeks after the injection. Scale bar, 5 cm. (F) Body weight of wild-type (+/+), *OT* KO (-/-), and *OT* cKO (*flox/-*) mice. The weight was measured at 5 weeks after injection of either vehicle or *AAV-Cre*. Note that this time point corresponds to 13 weeks old. **p < 0.01, one-way ANOVA with post hoc Tukey,s HSD. N = 10, 7, 9, and 10 for +/+, -/-, *flox/-* vehicle, and *flox/-* Cre, respectively. (G) Relationship between the number of remaining *OT*+ neurons in the PVH and the body weight of *OT^flox/-^* mice shown in (F). Magenta, exponential fit for the data from both Cre and vehicle. (H) Time course of daily food intake, defined as the average food intake in each week after *AAV-Cre* or vehicle injection. ***p < 0.001, Student’s t-test with post hoc Bonferroni correction. N = 6 each. (I) Cumulative food intake during the 5 weeks after the injection. ***p < 0.001, Student’s *t*-test. N = 6 each.

In addition to the PVH, OT neurons are also clustered in the SO (Zhang et al., 2021). To examine whether OT neurons in the SO also play inhibitory roles on body weight and food intake, we injected *AAV-Cre* into the bilateral SO of *OT^flox/-^* mice (Figure 2A and 2B). Similar to the PVH, the number of *OT*+ neurons in the SO significantly decreased at around 3 weeks after the injection (Figure 2C). Unlike the PVH, however, neither body weight nor food intake was significantly different compared with controls (Figure 2D–G), and no clear relationship was found between the number of the remaining **OT*+* neurons and body weight (Figure 2E). These results suggest that PVH OT neurons predominantly regulate food intake and body weight, and that SO OT neurons exert little influence.

**Figure 2.**
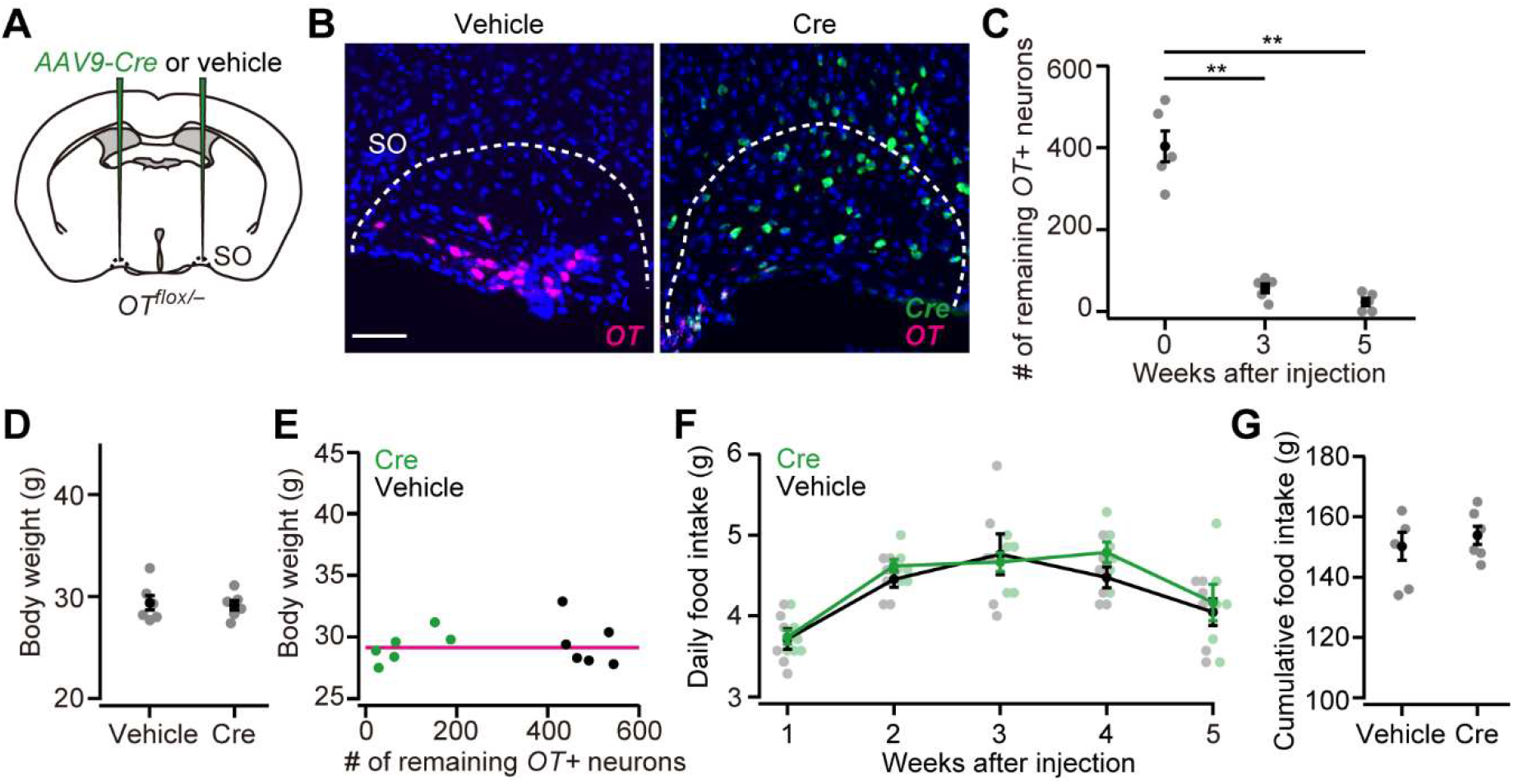
OT cKO in SO has a negligible effect on food intake and body weight. (A) Schematic of the virus injection. *AAV-Cre* or vehicle was injected into the bilateral SO of *OT^flox/-^* male mice. (B) Representative coronal sections of left SO from *OT^flox/-^* mice received vehicle (left) or *AAV-Cre* (right) injection. Five weeks after the injection. Magenta and green represent *OT* and *Cre in situ* stainings, respectively. Blue, DAPI. Scale bar, 50 μm. (C) The number of remaining *OT*+ neurons in the SO of mice that received *AAV-Cre* injection. **p < 0.01, one-way ANOVA with post hoc Tukey’s HSD. N = 5 each. (D) The body weight of *OT^flox/-^* mice did not differ between vehicle or *AAV-Cre* (Student’s t-test). N = 6 each. Data were obtained at 5 weeks after the injection. (E) Relationship between the number of remaining *OT*+ neurons in the SO and the body weight of *OT^flox/-^* mice shown in (D). Magenta, exponential fit for the data from both Cre and vehicle. (F) The time course of daily food intake was not statistically different (Student’s *t*-test with post hoc Bonferroni correction). N = 6 each. (G) Cumulative food intake during the 5 weeks after the injection. N = 6 each.

Because of the minor role of SO OT neurons, we focused on the PVH OT neurons in the following experiments.

### Weight of viscera and blood constituents

Increased food intake may influence not only body weight, but also the viscera and blood constituents. To examine these points, we collected internal organs and blood samples from non-fasted *OT^flox/-^* mice that had received either *AAV-Cre* or vehicle injection into the bilateral PVH (Figure 3A). While the weight of the stomach was unchanged (Figure 3B), a significant increase was observed in the weight of the liver in *OT^flox/-^* mice with *AAV-Cre* injection, likely because of the accumulation of fat in the liver (Figure 3C). We next measured the plasma concentration of glucose, triglyceride, and leptin. No significant differences in glucose levels were found (Figure 3D). In turn, the plasma concentrations of triglyceride and leptin were higher in *OT^flox/-^* mice that had received *AAV-Cre* injection than in those that had received vehicle injection (Figure 3D). Of note, a prominent increase in plasma leptin was also reported in the late-onset obesity cases of 6-month-old *OT* KO mice (Camerino, 2009). Our data regarding *OT* cKO showed the plasma leptin phenotype in the earlier stage of 13-week-old mice. These results suggest that the cKO of *OT* affects the homeostasis of viscera and blood constituents.

**Figure 3.**
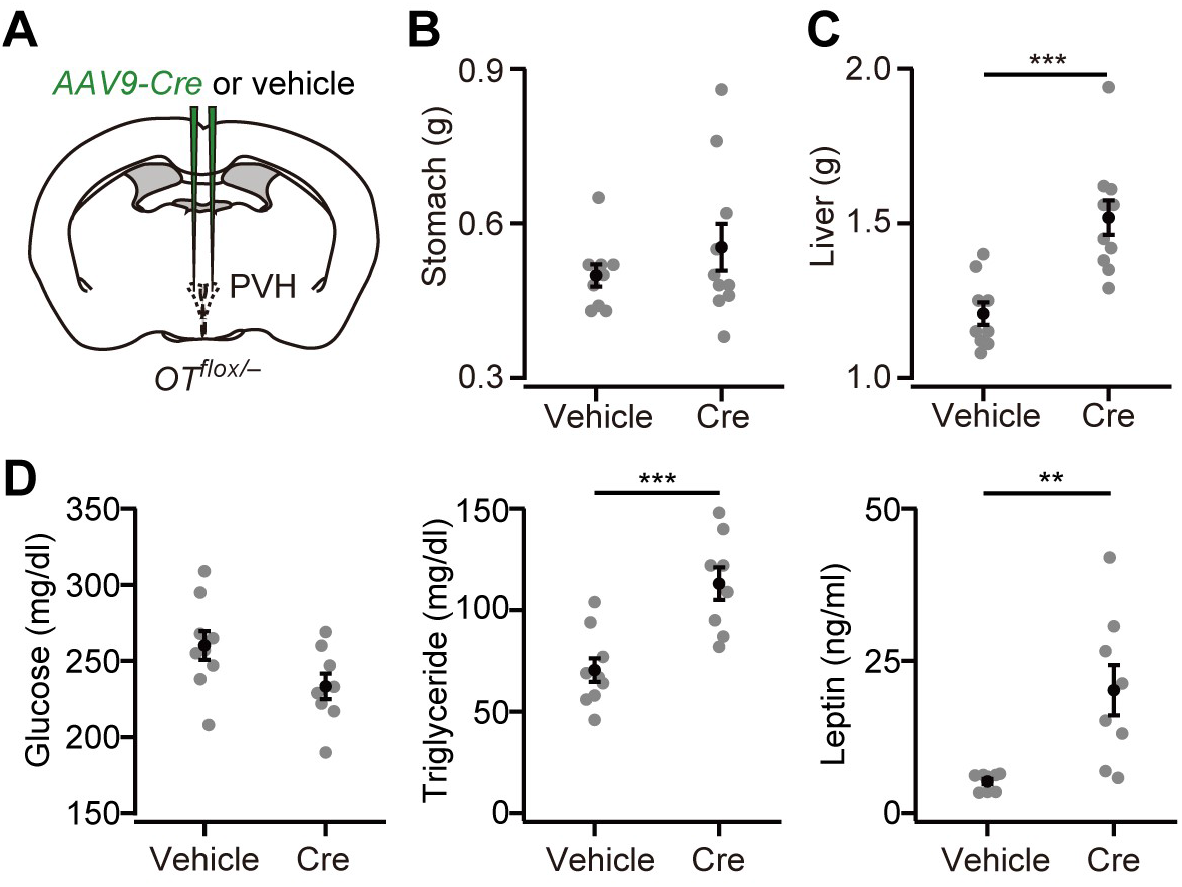
Weight of viscera and the blood constituents. (A) Schematic of the virus injection. *AAV-Cre* or vehicle was injected into the bilateral PVH of *OT^flox/-^* male. Data were obtained at 5 weeks after the injection. (B) The weight of the stomach was not statistically different (p > 0.5, Student’s t-test. N = 9 and 10 for vehicle and Cre, respectively). (C) The weight of the liver was significantly heavier in *AAV-Cre*-injected mice (***p < 0.001, Student’s t-test). (D) Plasma glucose (left), triglyceride (middle), and leptin (right) measured in the non-fasted *OT^flox/-^* mice. **p < 0.01, ***p < 0.001, Student’s *t*-test. N = 9 and 8 mice for vehicle and Cre, respectively.

### OT supplementation partially rescues *OT* cKO

If *OT* cKO caused increases in body weight and food intake with higher plasma triglyceride and leptin, such effects might be mitigated by the external administration of OT. This hypothesis is also supported by the fact that intraperitoneal (ip) or intracerebroventricular (icv) injection of OT has been shown to reduce both body weight and food intake (Maejima et al., 2018). To examine this hypothesis, first, we injected *AAV-Cre* into the bilateral PVH of *OT^flox/-^* mice, and from the next day of the injection, we conducted ip injection of OT (100 μL of 500 μM or 1 mM solution) once every 3 days (Figure 4A). Ip injection of the vehicle was used as control. At 5 weeks after the injection of AAV-Cre, OT-treated mice showed significantly reduced body weight, even though the number of remaining *OT*+ neurons was not significantly different (Figure 4B and 4C). We found that both daily food intake at 4–5 weeks and total food intake during the 5 weeks after the injection were also significantly reduced (Figure 4D and 4E). The reduction of body weight and food intake by our OT-treatment paradigm was somehow specific to *OT* cKO mice, given that neither reduction of food intake nor body weight was observed in wild-type males (Figure S2A–D). No significant improvement in the blood samples was found: both plasma triglyceride and leptin tended to be reduced in OT-treated mice, but did not reach the level of statistical significance (Figure 4F). These results suggest that external administration of OT can rescue at least the hyperphagic obesity phenotype of *OT* cKO.

**Figure 4.**
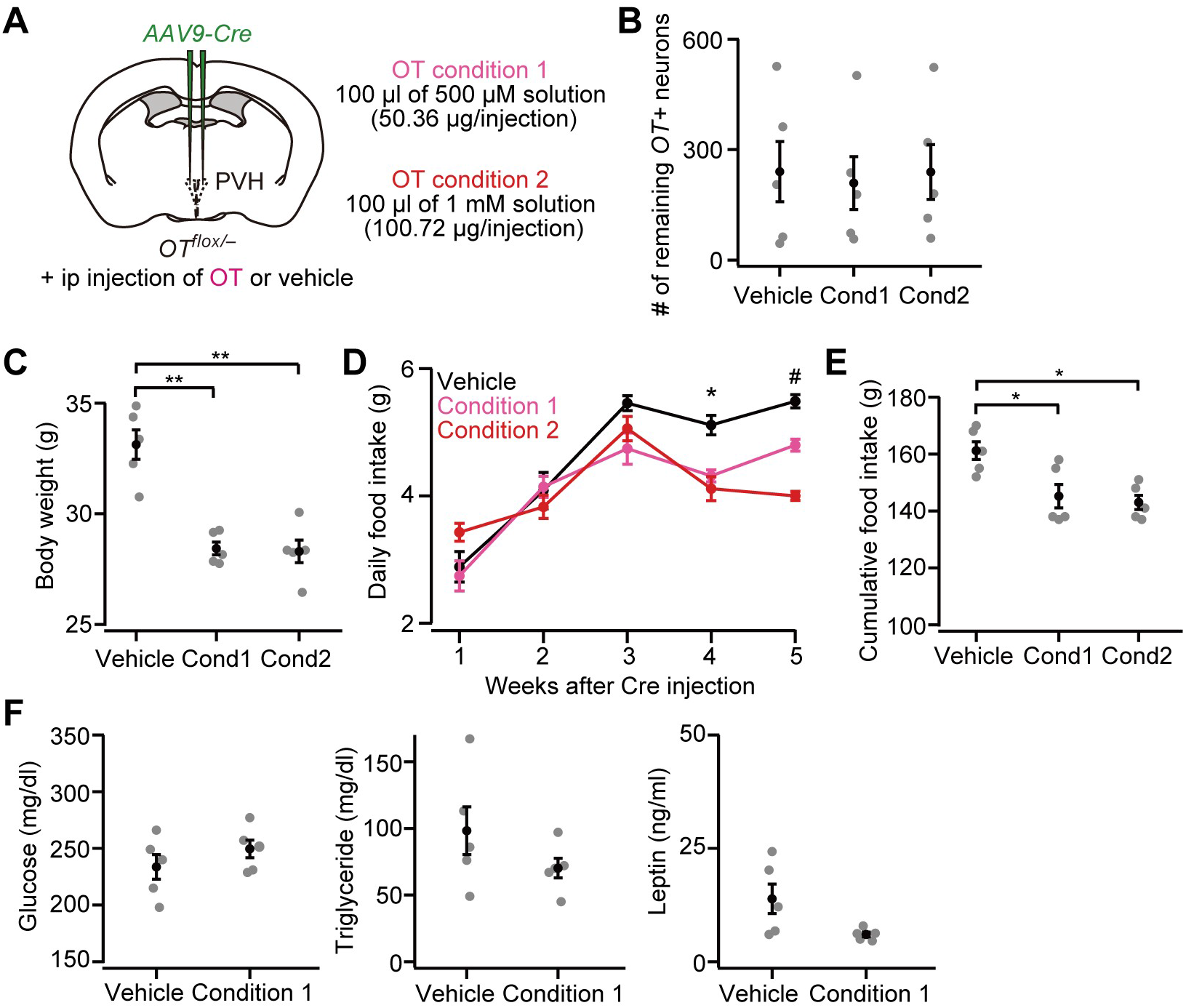
Ip injection of OT partially rescues PVH OT cKO phenotypes. (A) Schematic of the experiments. *AAV-Cre* was injected into the bilateral PVH of *OT^flox/-^* male mice. Data were obtained 5 weeks after the virus injection. Once every 3 days, the mice received ip injection of vehicle, 50.36 μg of OT (condition 1) or 100.72 μg of OT (condition 2) (see Materials and Methods). (B) The number of remaining *OT*+ neurons was not statistically different (p > 0.9, one-way ANOVA). Cond, condition. N = 5 each (same mice across panels B–E). (C) Ip injection of OT significantly decreased body weight. **p < 0.01, one-way ANOVA with post hoc Tukey’s HSD. Cond, condition. (D) Time course of daily food intake. Asterisks (*) denote significant differences for vehicle versus condition 1 and vehicle versus condition 2 (p < 0.05, Tukey’s HSD), and hashes (#) denote significant differences for vehicle versus condition 1, vehicle versus condition 2, and condition 1 versus condition 2 (p < 0.05, Tukey’s HSD). (E) Cumulative food intake during the 5 weeks after the virus injection was decreased in the mice that received ip injection of OT (*p < 0.05, one-way ANOVA with post hoc Tukey’s HSD). Cond, condition. (F) Plasma glucose (left), triglyceride (middle), and leptin (right) measured in the non-fasted *OT^flox/-^* mice. Decreases in triglyceride and leptin on average were found in OT-treated mice but did not reach the level of statistical significance (p = 0.314 and 0.065 for triglyceride and leptin, respectively, Student’s t-test). N = 5 each.

In addition to the daily food intake that we have examined so far, previous studies showed that mice ate less within several hours after receiving ip injection of OT (Arletti et al., 1989; Maejima et al., 2011). To examine whether *OT^flox/-^* mice that receive *AAV-Cre* injection similarly show reduced hourly food intake, we measured food intake after 6 h of fasting (Figure S2E and S2F). After fasting, the mice received an ip injection of OT, and food was provided again (Figure S2F). Cumulative food intake was measured at 1, 3, and 5 h after the placement of food (Figure S2F). Although the number of remaining *OT*+ neurons was comparable (Figure S2G), OT-injected mice ate less (Figure S2H). Taken together, ip injection of OT appears to reduce food intake on the scale of hours to days, thereby preventing the hyperphagic obesity induced by *OT* cKO in the PVH.

### OTR-expressing cells in the ARH mediate appetite suppression

Having established the importance of OT to suppress hyperphagic obesity, we examined the site of action of OT signaling that mediates appetite suppression. To this end, we prepared *OTR^flox/flox^* mice, in which the *OTR* gene can be knocked out under Cre expression (Takayanagi et al., 2005). Given that PVH OT neurons send dense projection to the hypothalamic nuclei (Yao et al., 2017; Zhang et al., 2021), and that OTR expression is also found in the hypothalamus (Fenselau et al., 2017; Mitre et al., 2016; Newmaster et al., 2020), we suspected that a fraction of appetite-suppression signals is mediated by the other nuclei of the hypothalamus. To test this possibility, we injected *AAV-Cre* (serotype 9) into the bilateral “anterior hypothalamus”, mainly aiming at nuclei such as the anteroventral periventricular nucleus (AVPV), medial preoptic nucleus medial part (MPNm), MPN lateral part (MPNl), and medial preoptic area (MPO) (Figure 5A; see Materials and Methods), and the “posterior hypothalamus”, containing nuclei such as the dorsomedial nucleus of the hypothalamus (DMH), ventromedial hypothalamic nucleus (VMH), lateral hypothalamic area (LHA), and ARH (Figure 5B). AAV-mediated *Cre* expression roughly covered these nuclei (Figure 5C and 5D). To examine if Cre expression reduced *OTR* expression, we visualized *OTR* mRNA using the RNAscope assay (see the Materials and Methods) (Sato et al., 2020; Wang et al., 2012). As *OTR* expression was observed as a dot-like structure (Figure 5E and 5F), we counted the number of such RNAscope dots in each DAPI+ cell. In a negative control experiment utilizing *OTR* KO mice, we often detected 1 or 2 RNAscope dots in the DAPI+ cells (Figure S3A and S3B). Therefore, we regarded a cell with three or more dots as an *OTR*-expressing cell (*OTR*+; Figure 5G, 5H, and S4A–C). We found that *AAV-Cre* injection successfully reduced the number of *OTR*+ cells in most of the targeted nuclei (Figure 5G and 5H). *OTR* cKO in the posterior but not anterior hypothalamus significantly increased body weight (Figure 5I and 5J). Similarly, a significant increase in food intake was observed in the mice that had received *AAV-Cre* injection into the posterior hypothalamus (Figure 5K and 5L).

**Figure 5.**
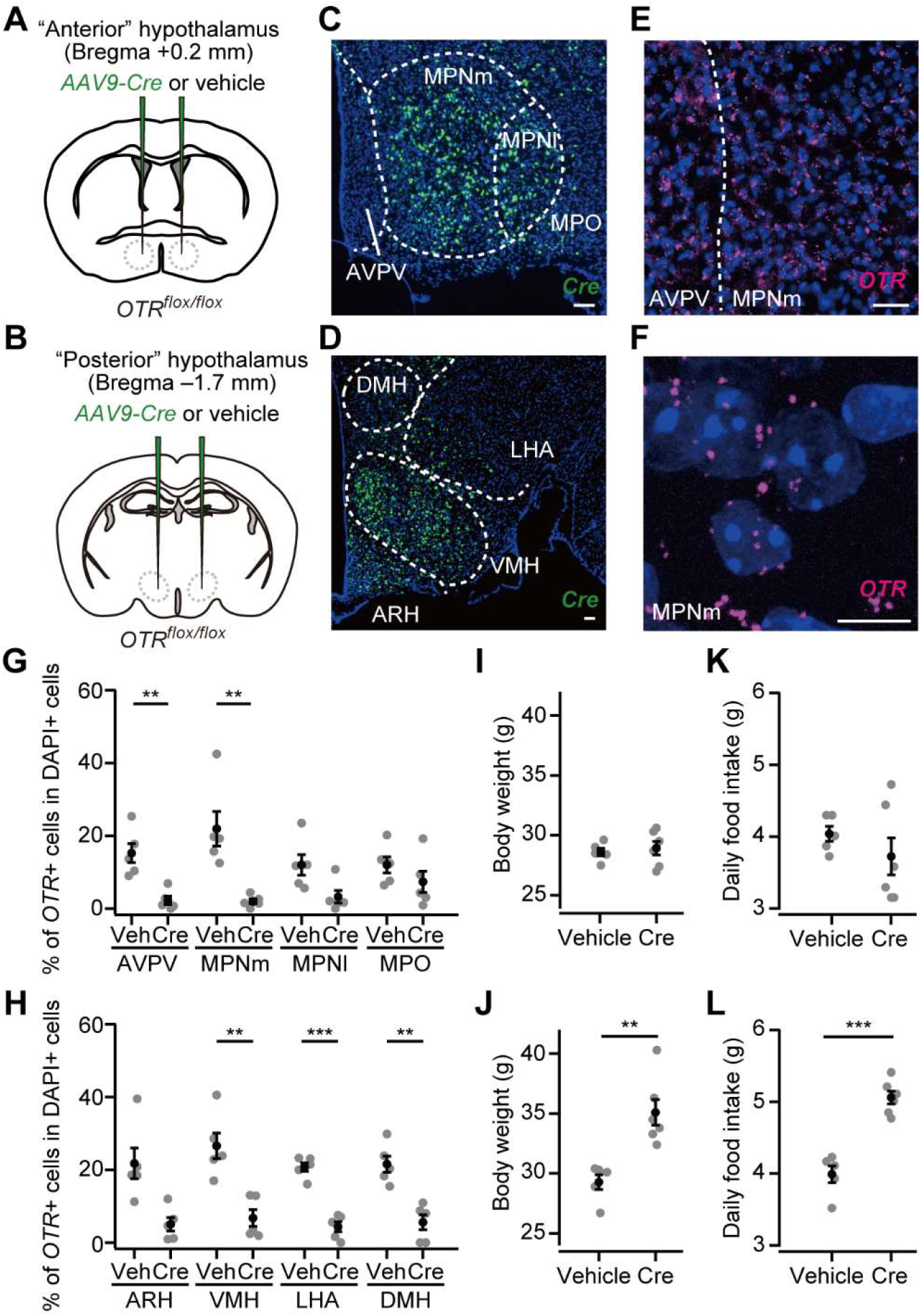
*OTR* cKO in the posterior hypothalamus induces increases in body weight and food intake. (A, B) Schematic of the virus injection. *AAV-Cre* or vehicle was injected into the bilateral anterior or posterior hypothalamus (see Materials and Methods) of *OTR^flox/flox^* male mice. (C, D) Representative coronal section showing *Cre* mRNA (green). Blue, DAPI. Scale bar, 50 μm. (E) A representative coronal section showing AVPV and MPNm from a vehicle-injected mouse. *OTR* mRNA was visualized by RNAscope (magenta). Blue, DAPI. Scale bar, 30 μm. (F) Projection of a confocal stack in MPNm from a vehicle-injected mouse. Magenta, *OTR* mRNA. Blue, DAPI. Scale bar, 5 μm. (G, H) Fraction of DAPI+ cells expressing OTR. Cells showing three or more RNAscope dots were defined as *OTR*+ (Figure S4). Veh, vehicle. N = 5 each. **p < 0.01, ***p < 0.001, Student’s t-test with Bonferroni correction. Decreases in the MPNl, MPO, and ARH on average were found in *AAV-Cre*-injected mice but did not reach the level of statistical significance in Student’s *t*-test with Bonferroni correction (p = 0.045, 0.289, and 0.012, respectively). (I, J) Body weight measured at 5 weeks after the injection. **p < 0.01, Student’s *t*-test. Anterior hypothalamus, N = 5 and 6 for vehicle and Cre, respectively, and posterior hypothalamus, N = 5 and 6 for vehicle and Cre, respectively. (K, L) Daily food intake measured at 5 weeks after the injection. ***p < 0.001, Student’s t-test. Anterior hypothalamus, N = 5 and 6 for vehicle and Cre, respectively, and posterior hypothalamus, N = 5 and 6 for vehicle and Cre, respectively.

We next aimed to pinpoint a specific nucleus in the posterior hypothalamus that could suppress hyperphagic obesity. To this end, we injected *AAV-Cre* (serotype 2) into the ARH or LHA (Figure 6A and 6B). This serotype of AAV-driven *Cre* expression was spatially localized (Figure 6C and 6D) compared with the AAV serotype 9 used in Figure 5. AAV-driven *Cre* expression reduced *OTR* expression in *OTR^flox/flox^* mice (Figure 6E and 6F). Body weight, daily food intake, and cumulative food intake were significantly greater in the mice that received *AAV-Cre* injection into the ARH, whereas no significant difference was found in the mice that expressed *Cre* in the LHA (Figure 6G–6L).

**Figure 6.**
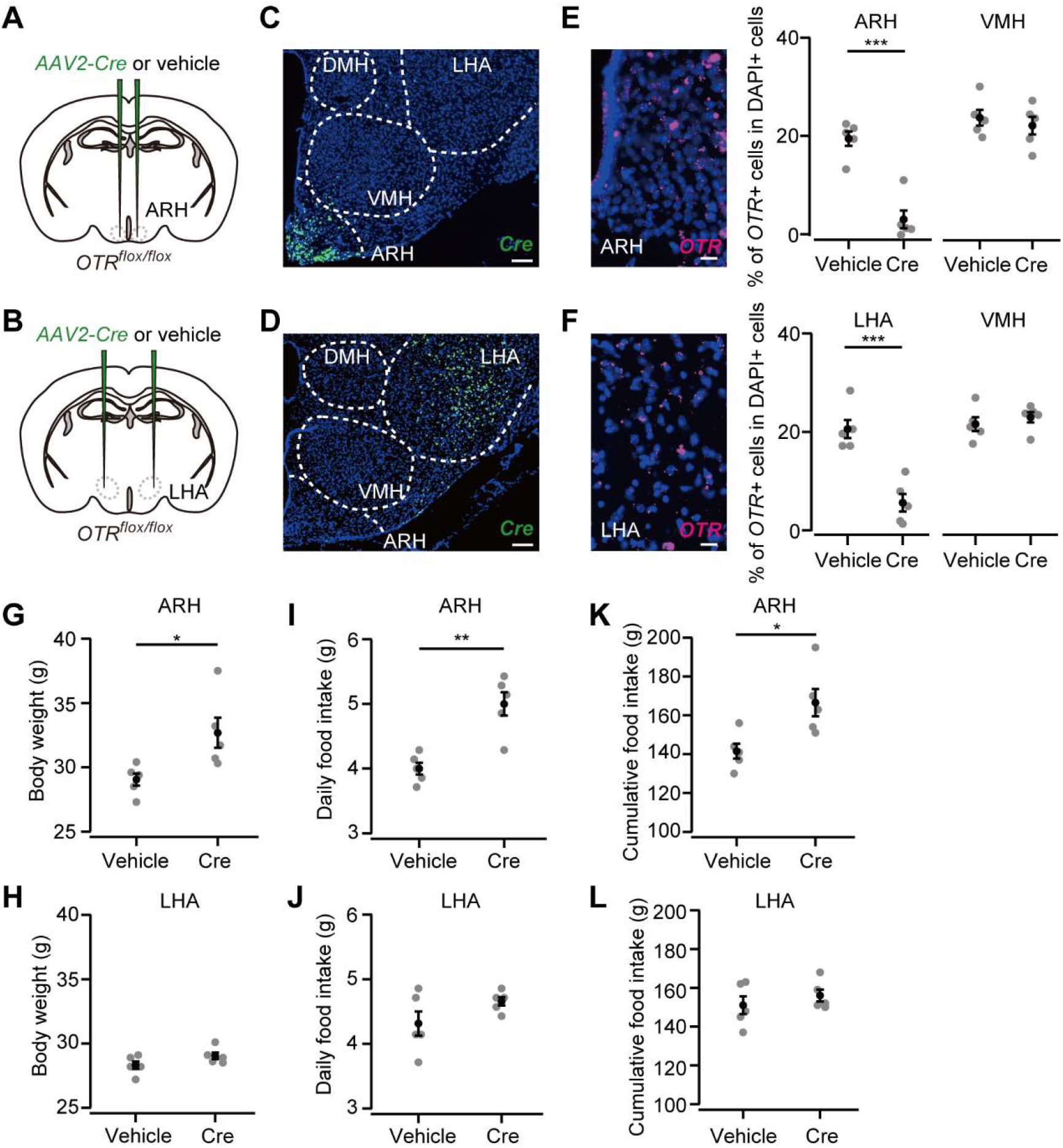
*OTR* expression in the ARH suppresses body weight and food intake. (A, B) Schematic of the virus injection. *AAV-Cre* or vehicle was injected into the bilateral ARH or LHA of *OTR^flox/flox^* male mice. (C, D) Representative coronal section showing *Cre* mRNA (green). Blue, DAPI. Scale bar, 50 μm. (E, F) Left, representative coronal section showing the ARH or LHA from a vehicle-injected mouse. *OTR* mRNA was visualized by RNAscope (magenta). Blue, DAPI. Scale bar, 5 μm. Right, fraction of DAPI+ cells expressing *OTR* in the ARH (E) or LHA (F) and VMH, a neighboring nucleus of the ARH and LHA. Cells showing three or more RNAscope dots were defined as *OTR*+. N = 5 each. ***p < 0.001, Student’s *t*-test with Bonferroni correction. (G, H) Body weight measured at 5 weeks after the injection. *p < 0.05, Student’s t-test. N = 5 each. (I, J) Daily food intake measured at 5 weeks after the injection. **p < 0.01, Student’s t-test. N = 5 each. (K, L) Cumulative food intake during the 5 weeks after the injection. *p < 0.05, Student’s t-test. N = 5 each.

Taken together, these results indicate that a fraction of the appetite-suppression signals from OT neurons is mediated by *OTR*-expressing cells in the posterior hypothalamus, especially those in the ARH.

## Discussion

### *OT* cKO increased body weight and food intake

In this study, we performed cKO of the *OT* gene by injecting *AAV-Cre*, which enabled region-specific KO of *OT*. By this advantage, we showed that OT produced by PVH OT neurons contributes to the regulation of body weight and food intake, whereas that by SO OT neurons does not (Figures 1 and 2). These data extend the previous results that mechanical disruption of PVH in rats increased both body weight and food intake (Shor-Posner et al., 1985; Sims and Lorden, 1986). In contrast to the *OT* cKO phenotype (Figure 1H and 1I), whole-body *OT* KO mice showed a normal amount of food intake (Camerino, 2009), suggesting compensational mechanisms. For example, when a certain gene is knocked out, expression of the related gene(s) is enhanced to compensate for some of the KO phenotypes functionally (El-Brolosy et al., 2019; Ma et al., 2019). A previous study showed elevated expression of dynorphin in the PVH of *OT* KO mice (Young et al., 1996). Given that dynorphin injection into the PVH promotes greater total caloric intake (Mattar et al., 2020), there may be an upregulation of genes that mitigate the excessive food intake caused by dynorphin, making the food intake of *OT* KO mice comparable to that of wild-type mice (Camerino, 2009). Transcriptomic analysis between *OT* KO and wild type may reveal a more complete picture of gene expression that can explain the compensational mechanisms in *OT* KO mice.

The phenotypic discrepancy between our data and diphtheria toxin-induced ablation of OT cells (Wu et al., 2012b) might be due to the loss of OT cells outside the PVH (maybe even outside the brain (Paiva et al., 2021)) that somehow elicited appetite suppression, and therefore counterbalanced the overeating phenotype caused by the loss of *OT* in the PVH. Alternatively, neuropeptides or neurotransmitters other than OT expressed in the PVH OT neurons might have conveyed appetite-stimulating signals, which remained intact in our *OT*-selective cKO model, but were disrupted in the cell-based ablation, resulting in only the overeating phenotype to appear in our case. Regardless of the scenario, our data establish the necessity of OT in the PVH to suppress overeating and suggest the presence of a hormone-based output pathway of PVH appetite regulation signals, in addition to the well-established neural pathways mediated by MC4R neurons (Garfield et al., 2015; Stachniak et al., 2014; Sutton et al., 2016).

### Downstream of OT neurons that mediate appetite-suppression signals

After eating a sufficient amount of food, animals stop eating owing to the appetite-suppression signals mediated in the brain. Several brain regions and cell types, such as Pomc-expressing neurons in the ARH (Ollmann et al., 1997; Sternson and Eiselt, 2017; Sutton et al., 2016), glutamatergic OTR-expressing neurons in the ARH (Fenselau et al., 2017), and calcitonin gene-related peptide expressing neurons in the parabrachial nucleus (Carter et al., 2013; Wu et al., 2012a), have been identified in this process. Our data showed that PVH OT neurons mediated appetite-suppression signals. Previous studies have shown that both oxytocinergic neurites and OTR-expressing neurons can be found in various brain and spinal cord regions (Jurek and Neumann, 2018; Lefevre et al., 2021; Newmaster et al., 2020; Oti et al., 2021; Zhang et al., 2021). In the present study, by *AAV-Cre* mediated cKO, we found that *OTR* expressed by neurons in the posterior hypothalamic regions, especially those in the ARH, mediates appetite-suppression signals (Figure 5 and 6). Our data are, therefore, generally consistent with the view that OTR-expressing neurons in the ARH evoke satiety signaling (Fenselau et al., 2017); however, we do not exclude the possibility that OTR in the other parts of the posterior hypothalamus, such as the VMH (Leng et al., 2008; Viskaitis et al., 2017) and medulla (Ong et al., 2017), also contributes to appetite suppression. Collectively, we suggest that one of the output pathways of the PVH for body weight homeostasis is mediated by OT signaling-based modulation of other hypothalamic appetite regulation systems.

We also showed that ip administration of OT can mitigate the overeating phenotype caused by the PVH *OT* cKO model (Figure 4). Together with the importance of OTR signaling in the ARH, one possibility is that the primary hypothalamic neurons that are located outside the blood–brain barrier (Yulyaningsih et al., 2017) directly receive ip-injected OT and transmit appetite-suppression signals. Alternatively, OTR-expressing neurons in the peripheral nervous system, such as those in the vagal sensory neurons transmitting intentional appetite-suppression signals (Bai et al., 2019), may indirectly modify feeding. Future studies should further dissect the responsible cell types and physiological functions of OTR signaling in the ARH. Recent advances in the real-time imaging of OTR activities, for example, with a circularly permuted green fluorescent protein binding to OTR (Ino et al., 2021; Qian et al., 2022), would be useful for delineating the circuit mechanism and spatiotemporal dynamics of the OT-mediated suppression of hyperphagic obesity, such as by pinpointing the site of OT release.

## Materials and Methods

### Animals

All experiments were conducted with virgin male mice. Animals were housed under a 12-h light/12-h dark cycle with *ad libitum* access to water and standard mouse pellets (MFG; Oriental Yeast, Shiga, Japan; 3.57 kcal/g). Wild-type C57BL/6J mice were purchased from Japan SLC (Hamamatsu, Japan). *OT* KO (Accession No. CDB0204E) and cKO (Accession No. CDB0116E) lines (listed at http://www2.clst.riken.jp/arg/mutant%20mice%20list.html) were generated and validated previously (Inada et al., 2022). The *OTR^flox/flox^* mouse line has been described (Takayanagi et al., 2005). *OTR* KO mice were generated by injecting *Cre* mRNA into the *OTR^flox/flox^* zygotes. We used only the mice that had the deletion allele without the *flox* allele by genotype PCR in the analysis (Figure S4). We also confirmed the result of Figure S4 in a small number of *OTR* KO mice that had been generated by conventional crossing from OTR *flox* mice. All animal procedures followed the animal care guidelines approved by the Institutional Animal Care and Use Committee of the RIKEN Kobe branch.

We chose the *OT flox/null* model to increase the efficiency of cKO (Figures 1–4). If we had used the *OT flox/flox* mice for cKO, because of high *OT* gene expression levels, a small fraction of the *flox* alleles that do not experience recombination would easily mask the phenotypes. Because *flox/null* alone (without Cre) has no phenotype (Figure 1F), we could justify the use of the *flox/null* model. Regarding *OTR* cKO (Figures 5 and 6), we chose the *flox/flox* model because the haploinsufficiency gene effect of OTR has been reported, at least in the context of social behaviors (Sala et al., 2013).

### Stereotactic viral injections

We obtained the AAV serotype 9 *hSyn-Cre* from Addgene (#105555; titer: 2.3 × 10^13^ genome particles/mL) and the AAV serotype 2 *CMV-Cre-GFP* from the University of North Carolina viral core (7.1 × 10^12^ genome particles/mL). To target the AAV or saline (vehicle) into a specific brain region, stereotactic coordinates were defined for each brain region based on the Allen Mouse Brain Atlas (Lein et al., 2007). Mice were anesthetized with 65 mg/kg ketamine (Daiichi Sankyo, Tokyo, Japan) and 13 mg/kg xylazine (X1251; Sigma-Aldrich) via ip injection and head-fixed to stereotactic equipment (Narishige, Tokyo, Japan). The following coordinates were used (in mm from the bregma for anteroposterior [AP], mediolateral [ML], and dorsoventral [DV]): PVH, AP –0.8, ML 0.2, DV 4.5; SO, AP –0.7, ML 1.2, DV 5.5; LHA, AP –2.0, ML 1.2, DV 5.2; ARH, AP –2.0, ML 0.2, DV 5.8. We defined the anterior and posterior hypothalamus by the following coordinates: anterior, AP +0.2, ML 0.2, DV 5.2; posterior, AP –1.7, ML 1.0, DV 5.2. The injected volume of AAV was 200 nL at a speed of 50 nL/min. After viral injection, the animal was returned to the home cage. In Figure 4, 100 μL of OT (1910, Tocris) dissolved in saline (vehicle) at 500 μM or 1 mM was ip-injected once every 3 days from the next day of *AAV-Cre* injection.

### Measurement of food intake

Food intake was measured by placing pre-weighted food pellets on the plate of a cage and reweighing them. In all the experiments, except those in Figure S2E–H, daily food intake was measured as follows: 200 g of food pellets were placed and food intake was measured once a week (weekly food intake). Daily food intake was calculated by dividing the weekly food intake by seven (days) and reported with significance digits of 0.1 g. In Figure S2E–H, after 6 h of fasting, 100 μL of OT dissolved in saline (vehicle) at 500 μM or 1 mM was ip-injected. Then, 80.0 g of food was placed and food intake was measured in units of 0.1 g after 1, 3, and 5 h.

### Fluorescent *in situ* hybridization

Fluorescent ISH was performed as previously described (Inada et al., 2022; Ishii et al., 2017). In brief, mice were anesthetized with sodium pentobarbital and perfused with PBS followed by 4% PFA in PBS. The brain was post-fixed with 4% PFA overnight. Twenty-micron coronal brain sections were made using a cryostat (Leica). The following primer sets were used in this study: *Cre* forward, CCAAGAAGAAGAGGAAGGTGTC; *Cre* reverse, ATCCCCAGAAATGCCAGATTAC; *OT* forward, AAGGTCGGTCTGGGCCGGAGA; and *OT* reverse, TAAGCCAAGCAGGCAGCAAGC. Fluoromount (K024; Diagnostic BioSystems) was used as a mounting medium. Brain images were acquired using an Olympus BX53 microscope equipped with a 10× (N.A. 0.4) objective lens. Cells were counted manually using the ImageJ Cell Counter plugin. In Figures 1C and 2C, cells were counted by an experimenter who was blind to the experimental conditions.

### RNAscope assay

*OTR* mRNA was visualized by the RNAscope Multiplex Fluorescent Reagent Kit (323110; Advance Cell Diagnostics [ACD]) according to the manufacturer’s instructions. In brief, 20-micron coronal brain sections were made using a cryostat (Leica). A probe against *OTR* (Mm-OXTR, 412171, ACD) was hybridized in a HybEZ Oven (ACD) for 2 h at 40 °C. Then, the sections were treated with TSA-plus Cyanine 3 (NEL744001KT; Akoya Biosciences; 1:1500). Fluoromount (K024; Diagnostic BioSystems) was used as a mounting medium. Images subjected to the analysis were acquired using an Olympus BX53 microscope equipped with a 10× (N.A. 0.40) or 20× (N.A. 0.75) objective lens, as shown in Figure 5E, 6E, and 6F. Figure 5F was obtained with a confocal microscope (LSM780, Zeiss) equipped with a 63× oil-immersion objective lens (N.A. 1.40). RNAscope dots were counted manually using the ImageJ Cell Counter plugin.

### Measurements of the weight of livers and stomachs

Mice were anesthetized with sodium pentobarbital. Livers and stomachs were obtained without perfusion and their weight was immediately measured. Before measurement, the stomach was gently pressed to eject the remaining contents.

### Plasma measurements

Mice were fed *ad libitum* before blood sampling. Mice were anesthetized with isoflurane and blood was collected from the heart. EDTA was used to prevent blood coagulation. Plasma concentrations of glucose, triglycerides, and leptin were measured by the enzymatic method, HK-G6PDH, and ELISA, respectively, through a service provided by Oriental Yeast (Shiga, Japan).

### Data analysis

All mean values are reported as the mean ± SEM. The statistical details of each experiment, including the statistical tests used, the exact value of n, and what n represents, are shown in each figure legend. The p-values are shown in each figure legend or panel; nonsignificant values are not noted. In Figure 1G and 2E, exponential fit was calculated by Igor (WaveMetrics).

## Author contributions

K.I. and K.M. conceived the experiments. K.I. performed the experiments, and K.I. and K.T. analyzed the data. M.Y. and K.N. provided the *OTR^flox/flox^* and *OTR* KO mice. K.I. and K.M. designed the experiments and wrote the paper.

## Acknowledgments

We wish to thank Mitsue Hagihara for the extensive technical support, the Laboratory for Comprehensive Bioimaging for the microscopy services, and the members of the Miyamichi lab for their comments on an earlier version of this manuscript. This work was supported by the RIKEN Special Postdoctoral Researchers Program, a grant from the Kao Foundation for Arts and Sciences, and JSPS KAKENHI (19J00403 and 19K16303) to K.I., and the JST CREST program (JPMJCR2021) and JSPS KAKENHI (20K20589) to K.M.

---

**Figure S1.**
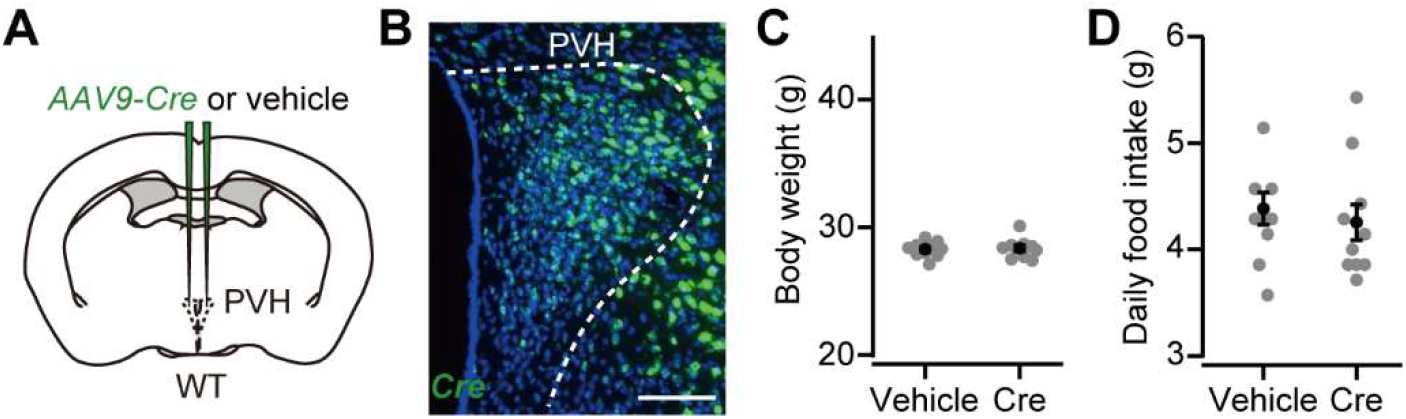
*AAV-Cre* injection into the PVH of wild-type mice does not induce hyperphagic obesity (related to Figure 1). (A) Schematic of the virus injection. *AAV-Cre* or vehicle was injected into the bilateral PVH of wild-type male mice. (B) Representative coronal section of the left PVH from wild-type mice that had received *AAV-Cre* injection. Data were obtained at 5 weeks after the injection. Green, *Cre in situ* staining, blue, DAPI. Scale bar, 50 μm. (C) Body weight measured at 5 weeks after the injection was not statistically different (p > 0.82, Student’s t-test). N = 10 each. (D) Daily food intake measured at 5 weeks after the injection (p > 0.59, Student’s *t*-test).

**Figure S2.**
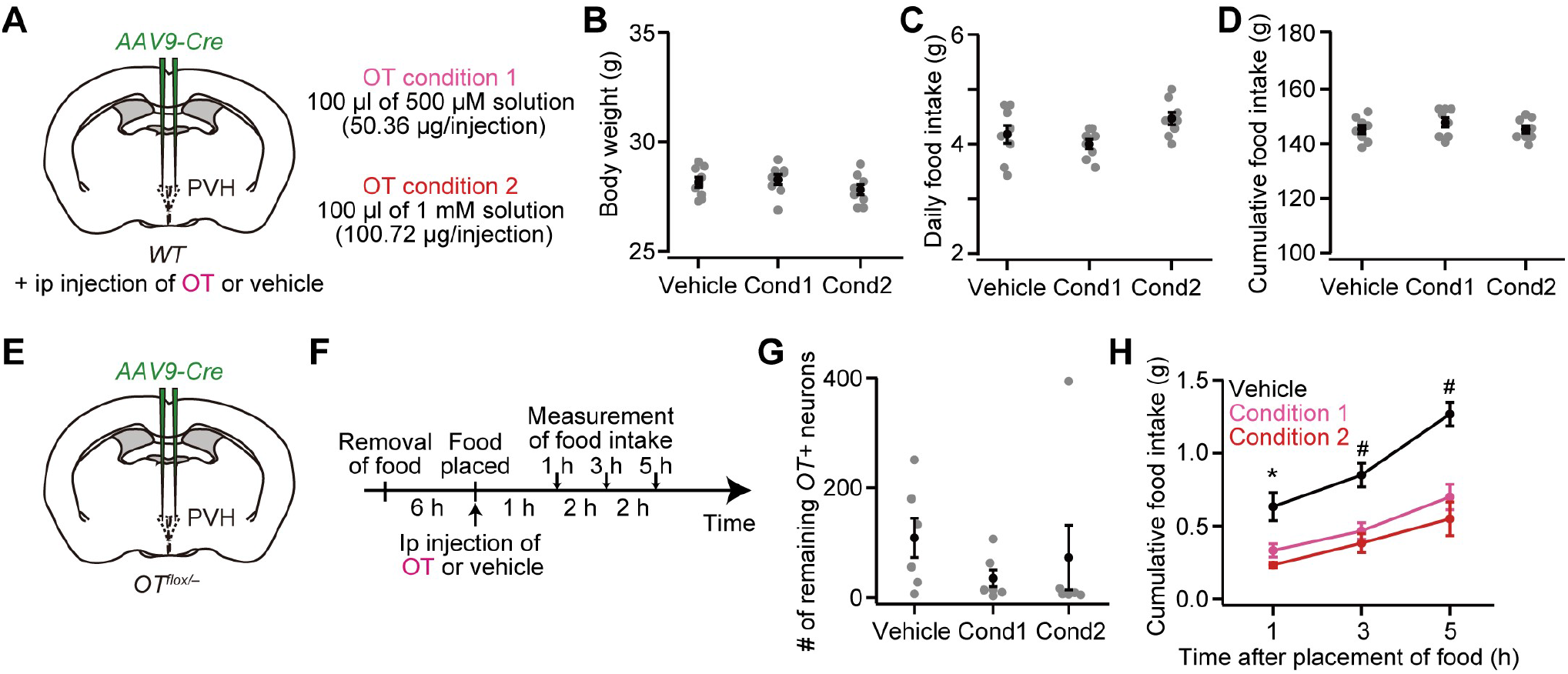
Ip injection of OT does not affect either body weight or food intake in wild-type mice, but reduces the hourly food intake of mice with OT cKO in the PVH (related to Figure 4). (A) Schematic of the experiments. *AAV-Cre* was injected into the bilateral PVH of wild-type male mice. The procedure used for the ip injection of OT solution was the same as that in Figure 4. (B) Body weight measured at 5 weeks after the injection was not statistically different (p > 0.41, one-way ANOVA). N = 8 each. (C) Daily food intake measured at 5 weeks after the injection (p > 0.068, one-way ANOVA). N = 8 each. (D) Cumulative food intake during the 5 weeks after the injection was not statistically different (p > 0.46, one-way ANOVA). N = 8 each. (E) Schematic of the experiments. *AAV-Cre* was injected into the bilateral PVH of *OT^flox/-^* male mice. Data were obtained at 5 weeks after the virus injection. (F) Schematic of the timeline of the experiment. Each mouse fasted for 6 h. Food intake was measured at 1, 3, and 5 h after the placement of food and ip injection of OT or vehicle. (G) The number of remaining **OT*+* neurons in the PVH was not statistically different (p > 0.52, one-way ANOVA). Cond, condition. N = 6 each (same mice across panels G and H). (H) Cumulative food intake. Asterisks (*) denote significant differences for vehicle versus condition 1 (p < 0.05, one-way ANOVA with post hoc Tukey’s HSD) and vehicle versus condition 2 (p < 0.01), and hashes (#) denote significant differences for vehicle versus condition 1 (p < 0.01) and vehicle versus condition 2 (p < 0.01).

**Figure S3.**
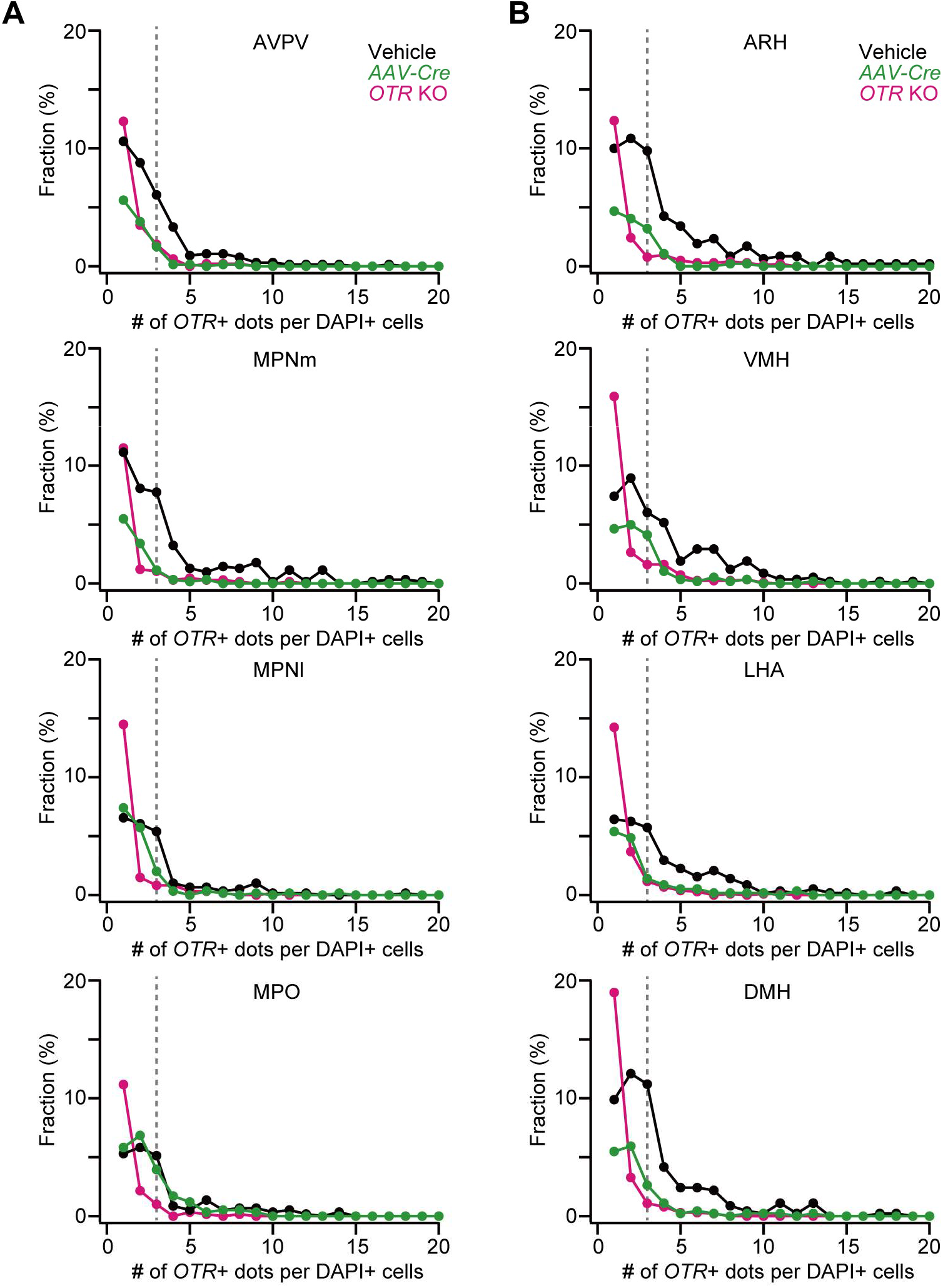
Distribution of *OTR*+ RNAscope dots, related to Figure 5. (A, B) Fraction of the number of *OTR*+ RNAscope dots in each DAPI+ cell in the nuclei in the anterior hypothalamus (A) or posterior hypothalamus (B). Black and green represent vehicle and *AAV-Cre* injected into the anterior hypothalamus (A) or posterior hypothalamus (B) in *OTR^flox/flox^* mice, respectively. Data from the same mice shown in Figure 5G and 5H. Magenta represents *OTR* KO mice (N = 3 and 5 for anterior and posterior, respectively). Dotted lines indicate three *OTR*+ dots in each DAPI+ cell.

**Figure S4.**
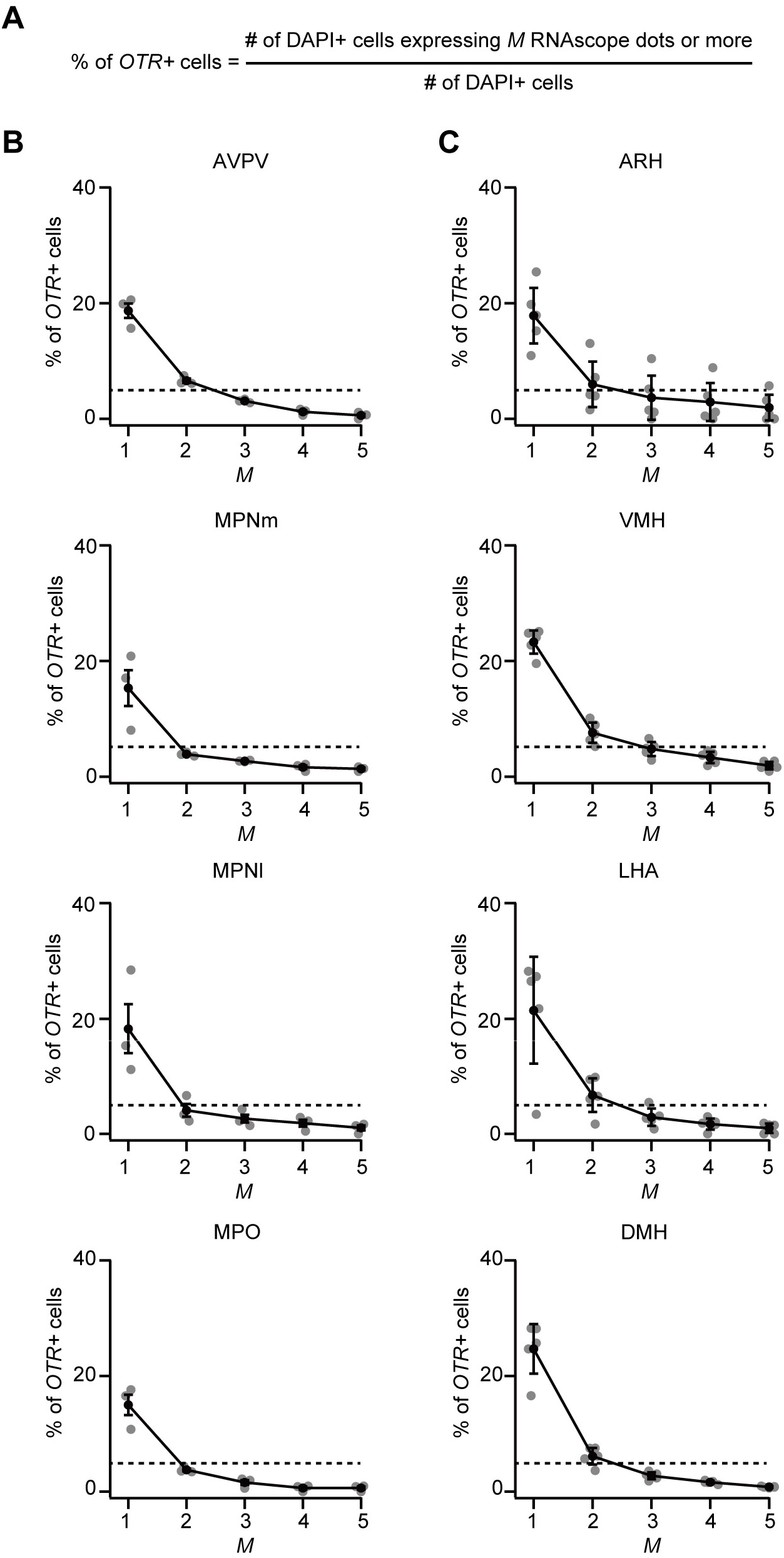
Testing the stringency in the definition of *OTR*+ cell by different numbers of RNAscope dots (related to Figure 5 and 6). (A) Calculation of the fraction of *OTR*+ cells as defined in this equation: we counted the number of RNAscope dots in each DAPI+ cell and defined a cell with *M* or more dots as an OTR-expressing cell in this figure. (B, C) Because we observed RNAscope dots in the OTR KO mice in which no OTR mRNA should be expressed, we evaluated the fraction of pseudo-positive detection with varying *M* values. Relationship between *M* and the fraction of *OTR*+ cells in the anterior hypothalamus (B) and posterior hypothalamus (C). Given that the pseudo-positive ratio on average was less than 5% (dotted line) in M ≥ 3, we defined a DAPI+ cell with three or more dots as an *OTR*+ cell in Figures 5 and 6. N = 3 and 5 mice for anterior and posterior, respectively.

## References

Andermann, M.L., and Lowell, B.B. (2017). Toward a Wiring Diagram Understanding of Appetite Control. Neuron 95, 757–778.

Arletti, R., Benelli, A., and Bertolini, A. (1989). Influence of oxytocin on feeding behavior in the rat. Peptides 10, 89–93.

Atasoy, D., Betley, J.N., Su, H.H., and Sternson, S.M. (2012). Deconstruction of a neural circuit for hunger. Nature 488, 172–177.

Bai, L., Mesgarzadeh, S., Ramesh, K.S., Huey, E.L., Liu, Y., Gray, L.A., Aitken, T.J., Chen, Y., Beutler, L.R., Ahn, J.S., et al. (2019). Genetic Identification of Vagal Sensory Neurons That Control Feeding. Cell 179, 1129–1143 e1123.

Balthasar, N., Dalgaard, L.T., Lee, C.E., Yu, J., Funahashi, H., Williams, T., Ferreira, M., Tang, V., McGovern, R.A., Kenny, C.D., et al. (2005). Divergence of melanocortin pathways in the control of food intake and energy expenditure. Cell 123, 493–505.

Blevins, J.E., Schwartz, M.W., and Baskin, D.G. (2004). Evidence that paraventricular nucleus oxytocin neurons link hypothalamic leptin action to caudal brain stem nuclei controlling meal size. Am J Physiol Regul Integr Comp Physiol 287, R87–96.

Camerino, C. (2009). Low sympathetic tone and obese phenotype in oxytocin-deficient mice. Obesity (Silver Spring) 17, 980–984.

Carter, M.E., Soden, M.E., Zweifel, L.S., and Palmiter, R.D. (2013). Genetic identification of a neural circuit that suppresses appetite. Nature 503, 111–114.

Catli, G., Acar, S., Cingoz, G., Rasulova, K., Yarim, A.K., Uzun, H., Kume, T., Kizildag, S., Dundar, B.N., and Abaci, A. (2021). Oxytocin receptor gene polymorphism and low serum oxytocin level are associated with hyperphagia and obesity in adolescents. Int J Obes (Lond) 45, 2064–2073.

Choi, S., Sparks, R., Clay, M., and Dallman, M.F. (1999). Rats with hypothalamic obesity are insensitive to central leptin injections. Endocrinology 140, 4426–4433.

Collaboration, N.C.D.R.F. (2016). Trends in adult body-mass index in 200 countries from 1975 to 2014: a pooled analysis of 1698 population-based measurement studies with 19.2 million participants. Lancet 387, 1377–1396.

Davis, C., Patte, K., Zai, C., and Kennedy, J.L. (2017). Polymorphisms of the oxytocin receptor gene and overeating: the intermediary role of endophenotypic risk factors. Nutr Diabetes 7, e279.

El-Brolosy, M.A., Kontarakis, Z., Rossi, A., Kuenne, C., Gunther, S., Fukuda, N., Kikhi, K., Boezio, G.L.M., Takacs, C.M., Lai, S.L., et al. (2019). Genetic compensation triggered by mutant mRNA degradation. Nature 568, 193–197.

Fenselau, H., Campbell, J.N., Verstegen, A.M., Madara, J.C., Xu, J., Shah, B.P., Resch, J.M., Yang, Z., Mandelblat-Cerf, Y., Livneh, Y., et al. (2017). A rapidly acting glutamatergic ARC->PVH satiety circuit postsynaptically regulated by alpha-MSH. Nat Neurosci 20, 42–51.

Garfield, A.S., Li, C., Madara, J.C., Shah, B.P., Webber, E., Steger, J.S., Campbell, J.N., Gavrilova, O., Lee, C.E., Olson, D.P., et al. (2015). A neural basis for melanocortin-4 receptor-regulated appetite. Nat Neurosci 18, 863–871.

Inada, K., Hagihara, M., Tsujimoto, K., Abe, T., Konno, A., Hirai, H., Kiyonari, H., and Miyamichi, K. (2022). Plasticity of neural connections underlying oxytocin-mediated parental behaviors of male mice. Neuron 110, 2009–2023.e5.

Ino, D., Hibino, H., and Nishiyama, M. (2021). A fluorescent sensor for the real-time measurement of extracellular oxytocin dynamics in the brain. bioRxiv, 2021.2007.2030.454450.

Ishii, K.K., Osakada, T., Mori, H., Miyasaka, N., Yoshihara, Y., Miyamichi, K., and Touhara, K. (2017). A Labeled-Line Neural Circuit for Pheromone-Mediated Sexual Behaviors in Mice. Neuron 95, 123–137 e128.

Jurek, B., and Neumann, I.D. (2018). The Oxytocin Receptor: From Intracellular Signaling to Behavior. Physiol Rev 98, 1805–1908.

Krashes, M.J., Shah, B.P., Madara, J.C., Olson, D.P., Strochlic, D.E., Garfield, A.S., Vong, L., Pei, H., Watabe-Uchida, M., Uchida, N., et al. (2014). An excitatory paraventricular nucleus to AgRP neuron circuit that drives hunger. Nature 507, 238–242.

Lawson, E.A., Marengi, D.A., DeSanti, R.L., Holmes, T.M., Schoenfeld, D.A., and Tolley, C.J. (2015). Oxytocin reduces caloric intake in men. Obesity (Silver Spring) 23, 950–956.

Lefevre, A., Benusiglio, D., Tang, Y., Krabichler, Q., Charlet, A., and Grinevich, V. (2021). Oxytocinergic Feedback Circuitries: An Anatomical Basis for Neuromodulation of Social Behaviors. Front Neural Circuits 15, 688234.

Lein, E.S., Hawrylycz, M.J., Ao, N., Ayres, M., Bensinger, A., Bernard, A., Boe, A.F., Boguski, M.S., Brockway, K.S., Byrnes, E.J., et al. (2007). Genome-wide atlas of gene expression in the adult mouse brain. Nature 445, 168–176.

Leng, G., and Sabatier, N. (2017). Oxytocin - The Sweet Hormone? Trends Endocrinol Metab 28, 365–376.

Leng, G., Onaka, T., Caquineau, C., Sabatier, N., Tobin, V.A., and Takayanagi, Y. (2008). Oxytocin and appetite. Prog Brain Res 170, 137–151.

Ma, Z., Zhu, P., Shi, H., Guo, L., Zhang, Q., Chen, Y., Chen, S., Zhang, Z., Peng, J., and Chen, J. (2019). PTC-bearing mRNA elicits a genetic compensation response via Upf3a and COMPASS components. Nature 568, 259–263.

Maejima, Y., Iwasaki, Y., Yamahara, Y., Kodaira, M., Sedbazar, U., and Yada, T. (2011). Peripheral oxytocin treatment ameliorates obesity by reducing food intake and visceral fat mass. Aging (Albany NY) 3, 1169–1177.

Maejima, Y., Yokota, S., Nishimori, K., and Shimomura, K. (2018). The Anorexigenic Neural Pathways of Oxytocin and Their Clinical Implication. Neuroendocrinology 107, 91–104.

Mattar, P., Uribe-Cerda, S., Pezoa, C., Guarnieri, T., Kotz, C.M., Teske, J.A., Morselli, E., and Perez-Leighton, C. (2020). Brain site-specific regulation of hedonic intake by orexin and DYN peptides: role of the PVN and obesity. Nutr Neurosci 22, 1105–1114.

Mitre, M., Marlin, B.J., Schiavo, J.K., Morina, E., Norden, S.E., Hackett, T.A., Aoki, C.J., Chao, M.V., and Froemke, R.C. (2016). A Distributed Network for Social Cognition Enriched for Oxytocin Receptors. J Neurosci 36, 2517–2535.

Newmaster, K.T., Nolan, Z.T., Chon, U., Vanselow, D.J., Weit, A.R., Tabbaa, M., Hidema, S., Nishimori, K., Hammock, E.A.D., and Kim, Y. (2020). Quantitative cellular-resolution map of the oxytocin receptor in postnatally developing mouse brains. Nat Commun 11, 1885.

Ollmann, M.M., Wilson, B.D., Yang, Y.K., Kerns, J.A., Chen, Y., Gantz, I., and Barsh, G.S. (1997). Antagonism of central melanocortin receptors in vitro and in vivo by agouti-related protein. Science 278, 135–138.

Onaka, T., and Takayanagi, Y. (2019). Role of oxytocin in the control of stress and food intake. J Neuroendocrinol 31, e12700.

Ong, Z.Y., Bongiorno, D.M., Hernando, M.A., and Grill, H.J. (2017). Effects of Endogenous Oxytocin Receptor Signaling in Nucleus Tractus Solitarius on Satiation-Mediated Feeding and Thermogenic Control in Male Rats. Endocrinology 158, 2826–2836.

Oti, T., Satoh, K., Uta, D., Nagafuchi, J., Tateishi, S., Ueda, R., Takanami, K., Young, L.J., Galione, A., Morris, J.F., et al. (2021). Oxytocin Influences Male Sexual Activity via Non-synaptic Axonal Release in the Spinal Cord. Curr Biol 31, 103–114 e105.

Paiva, L., Lozic, M., Allchorne, A., Grinevich, V., and Ludwig, M. (2021). Identification of peripheral oxytocin-expressing cells using systemically applied cell-type specific adeno-associated viral vector. J Neuroendocrinol 33, e12970.

Qian, T., Wang, H., Wang, P., Geng, L., Mei, L., Osakada, T., Tang, Y., Kania, A., Grinevich, V., Stoop, R., Lin, D., Luo, M., and Li, Y. (2022). Compartmental Neuropeptide Release Measured Using a New Oxytocin Sensor. bioRxiv, DOI: https://doi.org/10.1101/2022.02.10.480016

Sala, M., Braida, D., Donzelli, A., Martucci, R., Busnelli, M., Bulgheroni, E., Rubino, T., Parolaro, D., Nishimori, K., and Chini, B. (2013). Mice heterozygous for the oxytocin receptor gene (Oxtr(+/-)) show impaired social behaviour but not increased aggression or cognitive inflexibility: evidence of a selective haploinsufficiency gene effect. J Neuroendocrinol 25, 107–118.

Sato, K., Hamasaki, Y., Fukui, K., Ito, K., Miyamichi, K., Minami, M., and Amano, T. (2020). Amygdalohippocampal Area Neurons That Project to the Preoptic Area Mediate Infant-Directed Attack in Male Mice. J Neurosci 40, 3981–3994.

Shor-Posner, G., Azar, A.P., Insinga, S., and Leibowitz, S.F. (1985). Deficits in the control of food intake after hypothalamic paraventricular nucleus lesions. Physiol Behav 35, 883–890.

Sims, J.S., and Lorden, J.F. (1986). Effect of paraventricular nucleus lesions on body weight, food intake and insulin levels. Behav Brain Res 22, 265–281.

Stachniak, T.J., Ghosh, A., and Sternson, S.M. (2014). Chemogenetic synaptic silencing of neural circuits localizes a hypothalamus->midbrain pathway for feeding behavior. Neuron 82, 797–808.

Sternson, S.M., and Eiselt, A.K. (2017). Three Pillars for the Neural Control of Appetite. Annu Rev Physiol 79, 401–423.

Sutton, A.K., Myers, M.G., Jr., and Olson, D.P. (2016). The Role of PVH Circuits in Leptin Action and Energy Balance. Annu Rev Physiol 78, 207–221.

Takayanagi, Y., Kasahara, Y., Onaka, T., Takahashi, N., Kawada, T., and Nishimori, K. (2008). Oxytocin receptor-deficient mice developed late-onset obesity. Neuroreport 19, 951–955.

Takayanagi, Y., Yoshida, M., Bielsky, I.F., Ross, H.E., Kawamata, M., Onaka, T., Yanagisawa, T., Kimura, T., Matzuk, M.M., Young, L.J., et al. (2005). Pervasive social deficits, but normal parturition, in oxytocin receptor-deficient mice. Proc Natl Acad Sci U S A 102, 16096–16101.

Thienel, M., Fritsche, A., Heinrichs, M., Peter, A., Ewers, M., Lehnert, H., Born, J., and Hallschmid, M. (2016). Oxytocin’s inhibitory effect on food intake is stronger in obese than normal-weight men. Int J Obes (Lond) 40, 1707–1714.

Viskaitis, P., Irvine, E.E., Smith, M.A., Choudhury, A.I., Alvarez-Curto, E., Glegola, J.A., Hardy, D.G., Pedroni, S.M.A., Paiva Pessoa, M.R., Fernando, A.B.P., et al. (2017). Modulation of SF1 Neuron Activity Coordinately Regulates Both Feeding Behavior and Associated Emotional States. Cell Rep 21, 3559–3572.

Wang, F., Flanagan, J., Su, N., Wang, L.C., Bui, S., Nielson, A., Wu, X., Vo, H.T., Ma, X.J., and Luo, Y. (2012). RNAscope: a novel in situ RNA analysis platform for formalin-fixed, paraffin-embedded tissues. J Mol Diagn 14, 22–29.

Worth, A.A., and Luckman, S.M. (2021). Do oxytocin neurones affect feeding? J Neuroendocrinol, e13035.

Wu, Q., Clark, M.S., and Palmiter, R.D. (2012a). Deciphering a neuronal circuit that mediates appetite. Nature 483, 594–597.

Wu, Z., Xu, Y., Zhu, Y., Sutton, A.K., Zhao, R., Lowell, B.B., Olson, D.P., and Tong, Q. (2012b). An obligate role of oxytocin neurons in diet induced energy expenditure. PLoS One 7, e45167.

Yao, S., Bergan, J., Lanjuin, A., and Dulac, C. (2017). Oxytocin signaling in the medial amygdala is required for sex discrimination of social cues. Elife 6. e31373 DOI: 10.7554/eLife.31373

Young, W.S., 3rd, Shepard, E., Amico, J., Hennighausen, L., Wagner, K.U., LaMarca, M.E., McKinney, C., and Ginns, E.I. (1996). Deficiency in mouse oxytocin prevents milk ejection, but not fertility or parturition. J Neuroendocrinol 8, 847–853.

Yulyaningsih, E., Rudenko, I.A., Valdearcos, M., Dahlen, E., Vagena, E., Chan, A., Alvarez-Buylla, A., Vaisse, C., Koliwad, S.K., and Xu, A.W. (2017). Acute Lesioning and Rapid Repair of Hypothalamic Neurons outside the Blood-Brain Barrier. Cell Rep 19, 2257–2271.

Zhang, B., Qiu, L., Xiao, W., Ni, H., Chen, L., Wang, F., Mai, W., Wu, J., Bao, A., Hu, H., et al. (2021). Reconstruction of the Hypothalamo-Neurohypophysial System and Functional Dissection of Magnocellular Oxytocin Neurons in the Brain. Neuron 109, 331–346 e337.

